# Spatially-induced nestedness in a neutral model of phage-bacteria networks

**DOI:** 10.1101/125377

**Authors:** Sergi Valverde, Santiago F. Elena, Ricard Solé

**Affiliations:** ICREA-Complex Systems Lab, Universitat Pompeu Fabra - PRBB, Dr. Aiguader 88, 08003 Barcelona; Institut de Biologia Evolutiva (IBE), CSIC-UPF, Psg Maritim Barceloneta, 08003 Barcelona; Instituto de Biología Molecular y Celular de Plantas (IBMCP), CSIC-UPV, 46022 Valencia, Spain; Instituto de Biología Integrativa de Sistemas (I2SysBio), Consejo Superior de Investigaciones Científicas-Universitat de Valencia, 46182 Paterna, Valencia, Spain; Santa Fe Institute, 1399 Hyde Park Road, New Mexico 87501, USA

## Abstract

Ecological networks, both displaying mutualistic or antagonistic interactions, seem to share common structural traits: the presence of nestedness and modularity. A variety of model approaches and hypothesis have been formulated concerning the significance and implications of these properties. In phage-bacteria bipartite infection networks, nestedness seems to be the rule in many different contexts. Modeling the coevolution of a diverse virus-host ensemble is a difficult task, given the dimensionality and multi parametric nature of a standard continuous approximation. Here we show that this can actually be overcome by using a simple model of coevolving digital genomes on a spatial lattice and having exactly the same properties, *i.e*. a genome-independent fitness associated to fixed growth and death parameters. A matching allele model of phage-virus recognition rule is enough to generate a complex, diverse ecosystem with heterogeneous patterns of interaction and nestedness, provided that interactions take place under a spatially-constrained setting. It is found that nestedness seems to be an emergent property of the coevolutionary dynamics. Our results indicate that the enhanced diversity resulting from localized interactions strongly promotes the presence of nested infection matrices.

## I. Introduction

Our biosphere is a complex adaptive system where flows of energy and matter take place through tangled ecological networks (Montoya, Pimm, and Sole 2006). Most of these flows occur at the level of microorganisms, and microbial communities are in turn constantly coevolving with their viruses in highly dynamical ways. One dramatic illustration of the permanent arms race between bacteria and their viral partners is provided by the staggering scale of ecological interactions in marine ecosystems (Suttle 2005; Suttle 2007). It has been estimated that 10^23^ viruses might be present in the entire marine biotas, while no less than 10^23^ phage infections are taking place every second. The impact on population dynamics is not less impressive: bacteriophages might kill around 20% of the total microbial biomass in a single day. This massive turnover happens in an evolutionary context: bacteria and phages constantly (and rapidly) coevolve. Such coevolutionary arms races occur in all known examples, including the gut microbiome or soil ecosystems, and provide a source of both phenotypic and genotypic diversity while affecting community structure (Koskella and Brockhurst 2014).

On a large-scale perspective, the resulting networks of interaction between phages and their host microbes display a number of interesting regularities emerging from the underlying arms race dynamics (Weitz et al. 2013). One pervasive feature of the virus-host infection networks (along with modularity) is the presence of nestedness, namely the presence of a hierarchical pattern where (ideally) we can order both microorganisms and phages as illustrated in Figure 1. This type of pattern, which appears widespread in a wide array of contexts, has also been explained under rather different ways, from species-specific approaches grounded in the given community organisation to abstract statistical physics models. Some of these studies support the existence of optimisation principles that would pervade the nested architecture of ecological webs (Suweis et al. 2013). However, this idea has been challenged by further studies revealing that nested structures are likely to be an inevitable byproduct of other more fundamental properties of these graphs, in particular their heterogeneous character (Johnson, Dominguez-Garcia, and Munoz 2013; Feng and Takemoto 2014). What is the origin of nested webs in antagonistic systems?

The nested pattern found in phage-bacteria infection networks has been hypothesized to result from a coevolutionary sequence of adaptations driven by gene-for-gene recognition processes (Agrawal and Lively 2003; Flor 1956; Thompson and Burdon 1992; Weitz et al. 2013). In a dynamic gene-for-gene coevolutionary sequence, new mutations arising in the bacterial genome confer resistance to concurrent phages, while maintaining resistance to phages that were abundant in the past. Likewise, mutations in the concurrent phages result in their ability to infect these newly arose bacterial genotypes while still being able of infecting past bacterial genotypes (Bohannan and Lenski 2000). This process results in bacterial genotypes that are resistant to a subset of all possible phages and phages able of infecting a subset of all bacterial genotypes. In other words, the most infectious phage has access to most bacterial genotypes while the second most infectious phage has only access to a subset of these bacterial genotypes.

From the bacterial perspective, the most resistant bacteria can be only infected by a limited number of phages (usually the most infectious one) while the second most resistant bacteria can be infected a a larger number of phages (Figure 1). According to the gene-for-gene coevolutionary dynamics, fitness costs may appear to limit the phages to broader their host range without limits; bacteria also suffer of fitness costs that limit their capacity to resist all possible phages (Ashby and Boots 2017; Jover, Cortez, and Weitz 2013; Jover et al. 2015). The gene-for-gene model produces a wide variety of evolutionary outcomes that include stable genetic polymorphisms either within a range of infectivity or defence (Segarra 2005) or across multiple ranges provided direct frequency-dependent selection is on operation (Tellier and Brown 2007), and fluctuating selection between narrow and broad-range specialists and generalists (Agrawal and Lively 2003).

A popular alternative to the gene-for-gene model is the matching allele model (Agrawal and Lively 2003; Weitz et al. 2013). In this model, bacteria evolve resistance to a single phage genotype and lose resistance to other phages. Likewise, mutations in phage genomes confer the ability to infect new evolved bacterial genotypes while losing the capacity to infect ancestral bacterial genotypes. Henceforth, bacteria attempt to avoid the most common phage while phages seek to match the most common host (Frank 1993). The indirect negative frequency-dependent selection created by the matching allele model leads to fluctuating selection between equally highly specific genotypes. This high specificity between bacterial host and the viruses that can infect them translates into a modular structure in the phage bacteria infection networks (Weitz et al. 2013). In the extreme case in of a one-to-one matching between bacteria and phages, the resulting infection network is call monogamous (Korytowski and Smith 2015).

The majority of empirical evidences, some being generated within evolution experiments, from bacteria and phages (Bohannan and Lenski 2000; Flores et al. 2011), plants and diverse plant pathogens (Flor 1956; Hillung et al. 2014; Thompson and Burdon 1992), fruit flies and the sigma virus (Bangham et al. 2007), and fishes (Mouillot, Krasnov, and Poulin 2008; Vazquez et al. 2005) support that variation in hosts and parasites degrees of specialization is in general good agreement with the expectations from the gene-for-gene model. Modular infection networks have been also described at higher taxonomic levels, yet with a nested structure within each module (Flores, Valverde, and Weitz 2013; Roux et al. 2015).

In this paper we present a minimal model of bacteria-phage interaction that provides a minimal framework to address the problem of what are the requirements for evolving a nested infection network structure. Although other models have been formulated to that goal (Beckett and Williams 2013; Jover, Cortez, and Weitz 2013; Jover et al. 2015; Korytowski and Smith 2015) they rely on a large number of parameters and required some special assumptions concerning the shape of interaction functions. Here we have assumed the smallest amount of complexity by using a quasineutral model of bacteria-phage interactions that can account for the emergence of nested webs.

**Fig 1.**
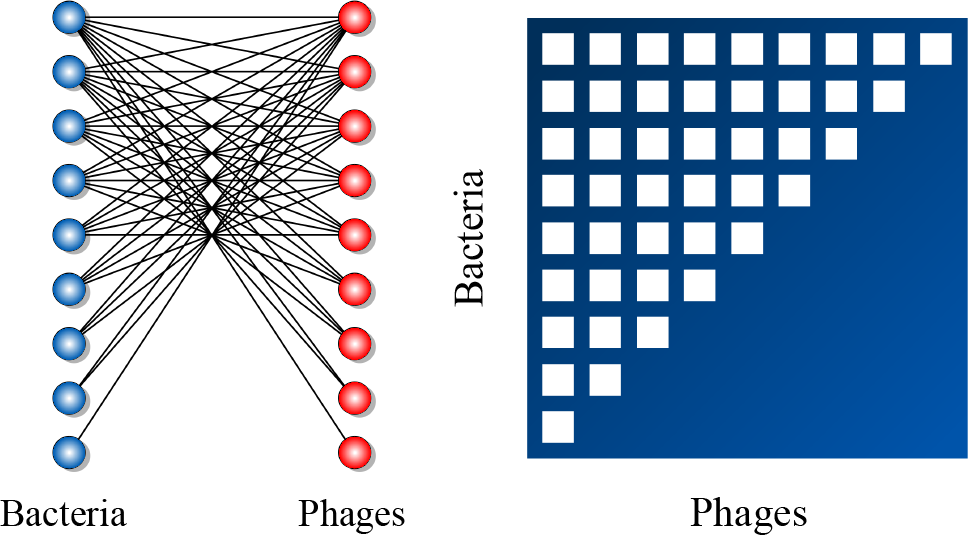
Many ecological networks are characterised by a pattern of nestedness. Our mathematical definitions of nestedness are derived from studies of bipartite networks (or two-mode) networks. A nested network displays a particular pattern of interactions that we can measure and detect. The left panel shows a bipartite network with two set of nodes. For example, one set corresponds to bacteria (blue balls) and the other to phages (red balls). Links represent interactions (infections) between pairs of dissimilar types. This bipartite network also accepts a matrix representation where rows and columns represent the two types of nodes and the entries of the matrix indicate the presence (white square) or absence (empty square) of pairwise interactions. The right panel shows an example of perfectly nested network, i.e., the non-zero elements of each row in the matrix are a subset of the non-zero elements in the subsequent rows.

## II. Neutral Coevolution Model

In this section we define our digital model of host-phage coevolution (all results published in this article are available upon request). Instead of using a model where a diverse repertoire of parameters is associated with each potential phenotype (resulting from a predefined genotype-phenotype mapping) we take the most simplifying assumption, namely a fully neutral system. In the spirit of other theoretical and modelling approaches (Alonso, Etienne, and McKane 2006) the species-level idiosyncrasies are ignored in favour of an upper-level of description. By using this toy model approach, we hope to gather insight into the network-level universals.

The model considers two populations of replicators, namely phages and bacteria, each represented as a *v*–dimensional string. Specifically, we indicate as 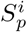 and 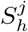, respectively the digital genomes (bit strings) associated to the *i*-th and *j*-th phage and bacteria genotypes. In other words, we have the strings given by the bit sequences:

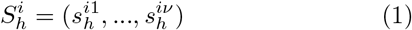

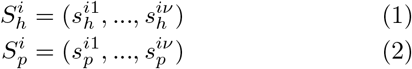

with 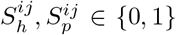 where *i* = 1,.…,2^*v*^ Both populations reproduce and evolve on a two-dimensional space with toroidal boundary conditions. In Figure 2 we display the structure of the coupled interaction between phages and bacteria, which will interact under a genomematching rule, defined below.

**Fig 2.**
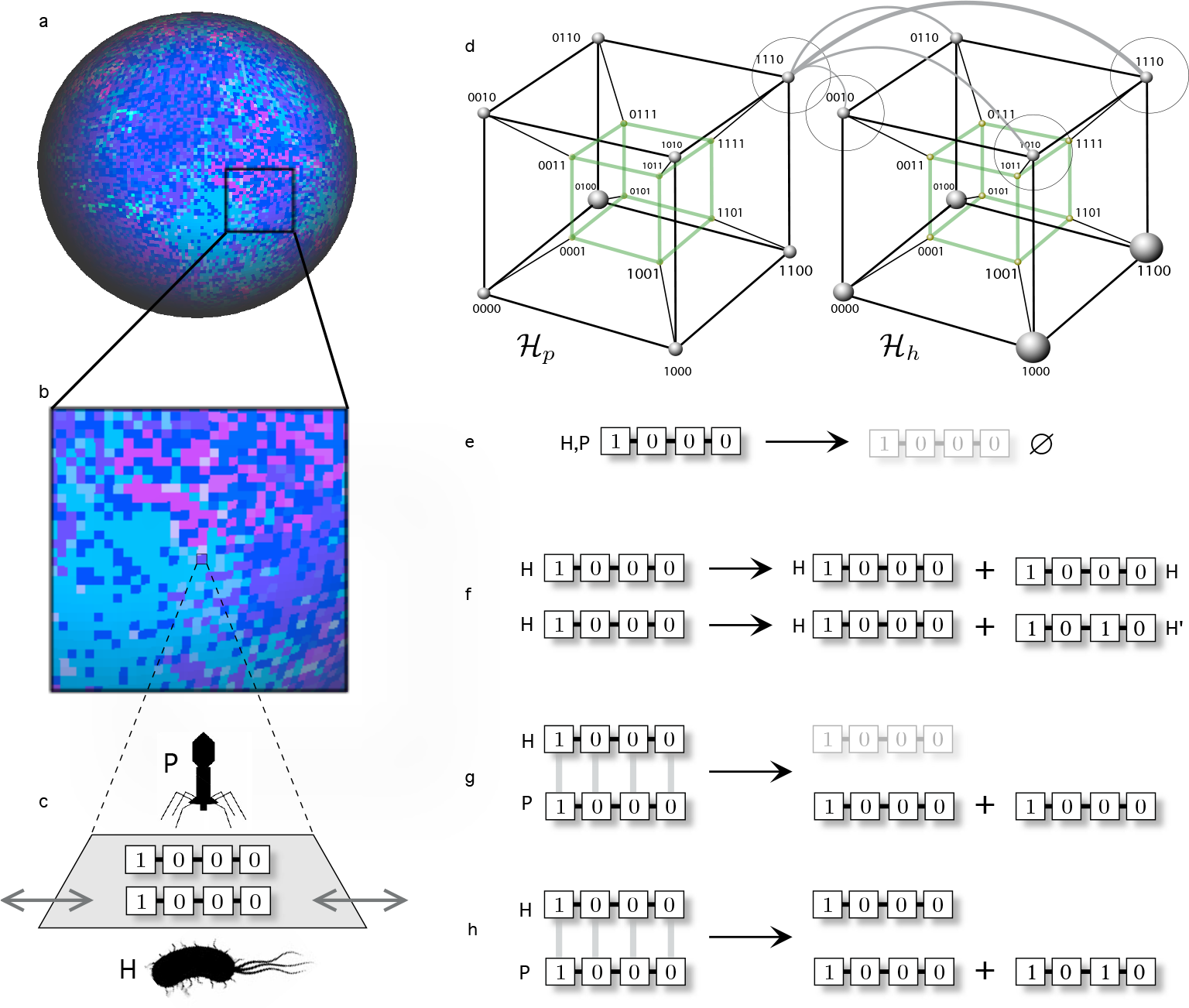
Coevolution in coupled landscapes associated to a phage-bacteria model based on a digital genomes representation. Each individual is described by means of a digital genome of length *v*. The model dynamics is defined on a two-dimensional surface (a) where the color of each site indicates (in this case) the presence of different strings. In (b) an expanded area show the presence of local patchiness where each site (c) can be occupied by one string of each class. Strings can move randomly to neighbouring free sites, as indicated by the grey arrows. From the point of view of the genotype space, we have two coupled sequence hypercubes (here *v* = 4) where similarity between phage and bacteria recognition sequences (*i.e*. the matching allele model or recognition, determines the probability of interaction. Two different hypercubes are shown, one for phages (left) and another (right) for bacteria. One given virus (like 1110) will be able to interact with those bacteria whose genome is closer (the probability of this interaction is indicated by weighted gray lines). The basic rules used in the model are summarised in (e–h). These are: (e) sequence removal (death), (f) replication of host string, which can be accurate *H → 2H* or inaccurate*H → H + H′*, for the phage-bacteria interaction.

The model is intended to define a minimal setting of rules including a random death of strings at a given rate, which can be represented as decay reactions, namely
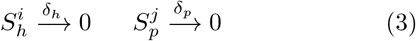
independent on their specific genome sequence. The bacterial strains are assumed to replicate leading to two identical copies with a probability that depends on the mutation rate, namely 
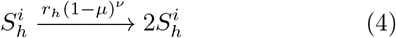
where(1-µ)*^v^*is the probability that all the bits are properly copied (no mistakes occur). Any mutation in at least one bit in the string will lead to a different sequence, namely 
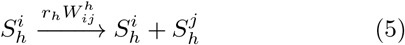
where 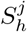 will be usually a one-mutation neighbour in sequence space (provided that mutation rates are small enough) but in general the probability associated to a mutation from 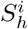 to 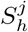 will be 
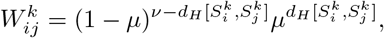
being 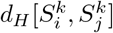 the Hamming distance between two sequences:

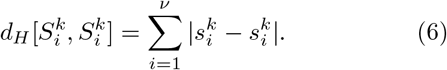

**Fig 3.**
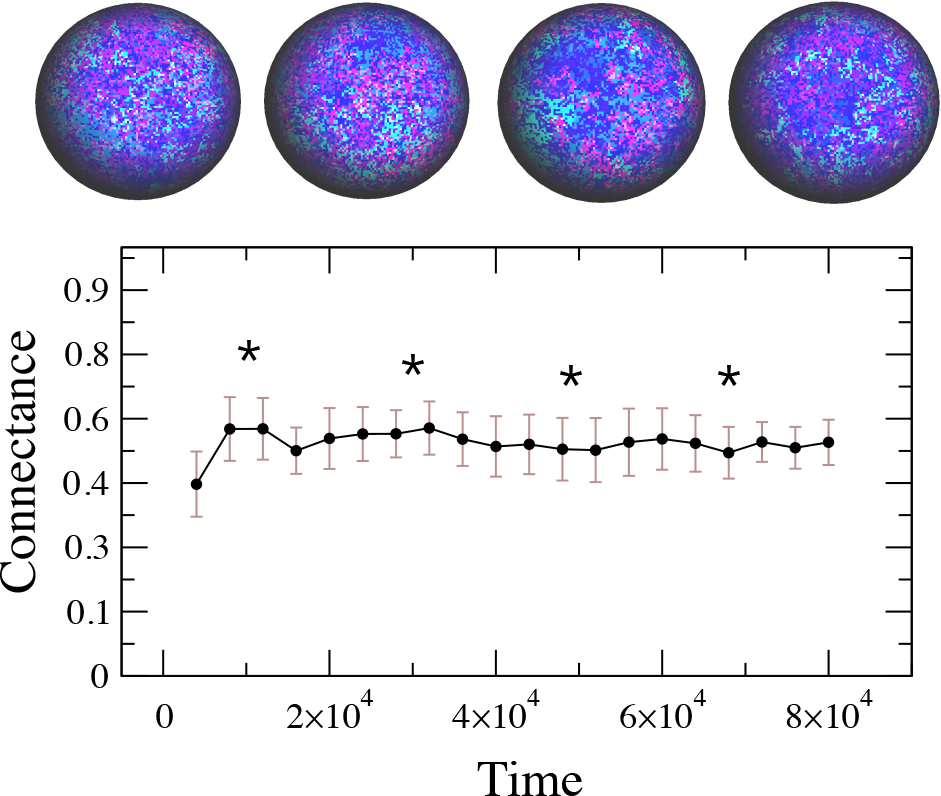
Temporal dynamics of connectance in a typical run of the model. The connectance stabilises around a well-difined *C* ≈ 0.45.This density of links allows for a high diversity of possible system configurations. From left to right, the snapshots (whose location is pointed with stars in the curve) display an evolving pattern of spatial heterogeneity. Here we use the same parameters described in the main text.

The reproduction of the phage requires the infection of a bacterial cell provided that a genome matching occurs. If the matching is perfect (and thus *d_H_* = 0) we assume that the interaction occurs with probability *ϕ* = 1. but if *d_H_* > 0 the probability *ϕ* will decay linearly with the Hamming distance (since recognition and matching are less accurate):

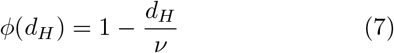

Specifically, two strings belonging to a phage 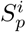 and a host 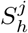 will lead to an error-free reaction:

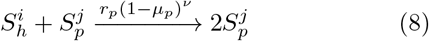
and the alternative scenario with a mutated offspring:

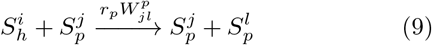
where the term 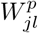 is defined as before.

The final set of rules involves the spatial dynamics of strings on the lattice. Each site in this lattice can be occupied by one string of each class. Host and parasite strings move, independently and randomly, to empty neighbour cells with diffusion probabilities *D_h_* and *D_p_* respectively. To simplify the analysis, all the simulations were run with maximum diffusion constants *D_h_ = D_p_* = 1, also setting *δ_h_ = δ_p_* = 10^−2^. In this way, we strongly reduce the parameter space, considering the diffusion and decay properties of all strings identical, no matter their precise sequence.

The rest of the parameters has been chosen as follows. The mutation rate of the virus must be larger than the one exhibited by the host. Here we use a mutation rate *µ* = 10^−4^ for the phage and *µ* = 10^−5^ for the bacteria. Other parameters have been used (avoiding very high rates that can lead to an error catastrophe) and similar results of those reported here have been found. Finally, the replication parameter *r* of the host strings is fixed to *r* = 0.8 An important point to be made here is that our genome-independent parameters mades our model a highly homogeneous one, that is an effectively neutral model, except for the functional dependence associated to the matching rule. The lattice was randomly inoculated by either bacteria and phage random sequences, starting from an initial condition were 25% of sites are randomly seeded by phages and the same amount (but different random sites) for the bacteria. All initial strings are identical, defined by the sequence 1000. As we can see from this model description, the whole dynamics will lead (if both populations are present and parameters allows) to an arms race that is limited to a constant movements through the sequence hypercube.

## III. Spatial Dynamics And The Emergence Of Bacteria-Phage Bipartite Nested Networks

The degree of interaction among species is not randomly distributed and captures different ecological and evolutionary factors. Disentangling these components requires a combination of empirical measurements and of theoretical models (see below). How ecological, genetics and epidemiological processes interact to generate and maintain structural variation? What is the general structure of bacteria-virus infection networks? Empirical and theoretical studies have shown that bipartite networks can be (i) nested, *i.e*., the interactions between nodes can be represented as subsets of each other (Flores et al. 2011), (ii) modular, *i.e*., a network composed of densely connected groups of nodes, and (iii) multi-scale, *i.e*., the network shows different features depending on whether the whole or smaller components are under consideration (Flores, Valverde, and Weitz 2013). In this study, we are particularly interested in how the spatial component influences the emergence of nestedness (see next section).

An infection network involves two disjoint subsets of species, *i.e*., bacterial hosts and viruses. Any pair of species is always related (*i, j*) ∈ *E* provided that pathogen *j* can infect host *i*. The set of links of this network can be described with the (binary) adjacency matrix *A* = [*A_i, j_*] in which *A_i, j_* = 1(presence) if the nodes *i* and *j* are connected or *A_i, j_* = 0 (absence), otherwise.

The degree of a species
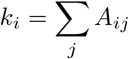
is the number of connections attached to this node. Now, assume that hosts are indexed 1, 2, …, *N_H_* and viruses are labeled *N_H_* + 1, *N_H_* + 2, …, *N_H_* + *N_V_* where *N_H_* is the number of bacterial species, *N_V_* is the number of virus species, and *N* = *N_H_* + *N_V_* is the total number of species in our system. Using this vertex labelling approach, we can show the adjacency matrix has a block off-diagonal form as follows:

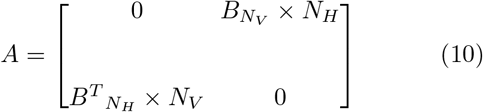
where *B* is the *N_H_* × *N_v_* incidence matrix, 0 is the allzero matrix that reflects the bipartite constraint, *i.e*., we only allow interactions between alike species.

**Fig 4.**
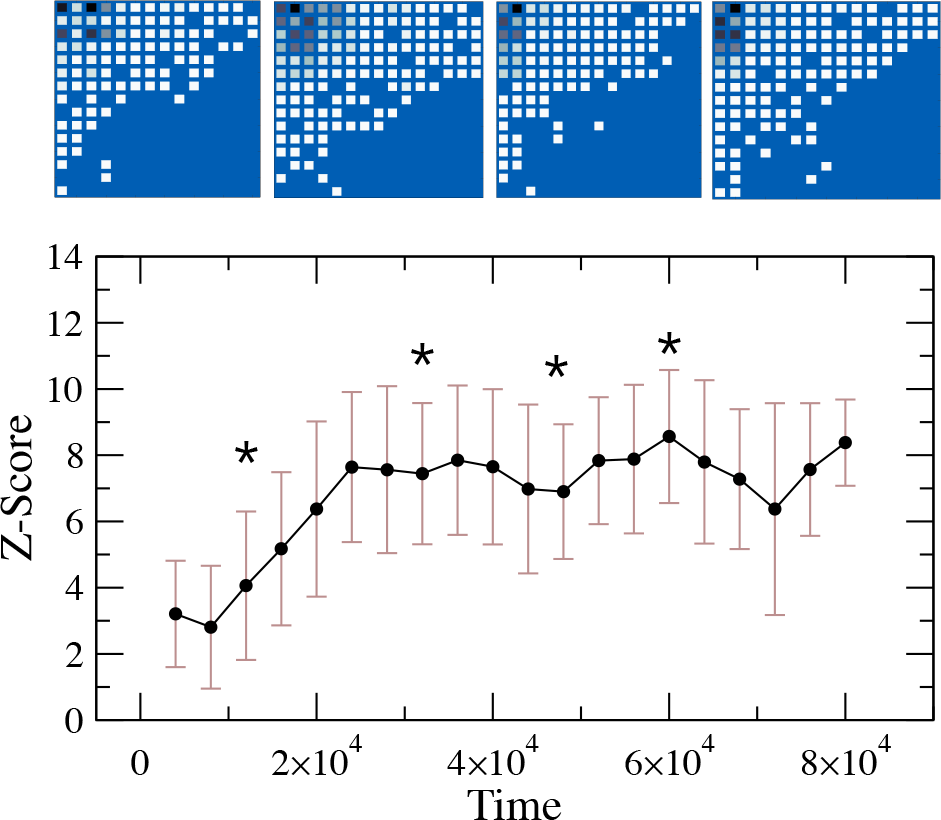
Temporal dynamics of the statistical significance of nestedness. After an initial transient period, the global organisation of the *γ*-matrix settles in stable nested patterns (the average *Z* ≈ 5). The top row shows several snapshots of the bacteria-phage interaction network taken at different evolutionary stages (whose location is given by stars in the curve). Both the binary structure and the quantitative preference matrix are significantly nested. The matrix at time *t* has been obtained by first counting the frequency of interactions *B_i, j_* observed in the time period [*t* − Δ*t, t*] (here Δ*t* = 2000 time steps) and then discarding the mass action term (*x_i_x_j_*). In each matrix, darker colours represent higher interaction preferences. Tests for nestedness are based in the null model described in the main text.

Reliable nestedness measurement takes into account the strength of infections in the bipartite network. For example, previous studies have shown that ubiquitous nestedness of binary adjacency matrices (Flores et al. 2011) is not always reproduced by quantitative studies (Staniczenko, Kopp, and Allesina 2013). The entries of a quantitative incidence matrix *B* take values different from 1 or 0, like the number of infected individuals. In both the binary and quantitative cases, an incidence matrix is perfectly nested when its rows and columns can be sorted such that

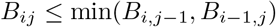

with *B*_1,*j*_ > 0 and *B*_*i*, 1_ > 0 for all 1 ≤ *i ≤ N_P_* and 1 ≤ *j* ≤ *N_V_*, that is, the set of edges in each row *i* contains all the edges in row *i* + 1 while the set of edges in column *j* contains the set of edges in column *j* + 1.

The above suggests a costly approach to maximal nestedness that looks for the optimal matrix ordering. This algorithm has several computational disavantages and it is, in fact, the basis for many published methods. The spectral radius *ρ*(*B*)(or the largest eigenvalue of the matrix) gives a natural scale for nestedness, with higher spectral radius corresponding to more nested configurations (Staniczenko, Kopp, and Allesina 2013). The spectral radius of the incidence matrix has two useful properties: (i) matrix eigenvalues are independent of arbitrary permutations of rows and columns and (ii) this quantity can be derived for both binary and quantitative infection networks.

We now investigate the temporal evolution of nestedness in our model. Nestedness is a relative value that depends on the size and the density of interactions. In our model, the number of species *N* it is bounded by the fixed genome lengths. On the other hand, link density is a key parameter defined by the network connectance *C*, or the proportion of possible network connections i.e. *C = L/N*^2^, where the number of links *K* is simply 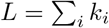.

It has been proposed that the stability of dynamical processes constraints the possible values of connectance (May 1972). In our computational simulations, we have observed that connectance reaches an average value *C* ≈ 0.45 (see Figure 3). A detailed analysis of interactions reveal a highly dynamical system where connections among species are constantly added or removed while keeping the same average connectance.

In general, there is a contribution 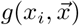 to the growth of population *i* from interactions with other species in the system 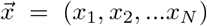 where *x_i_* is the population density of individual species (Staniczenko, Kopp, and Allesina 2013). We can further divide the interaction between any pair of species (*i, j*)in two components: the frequency of interactions *γ_i, j_x_i_x_j_* and the effect of each interaction 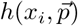. Then,

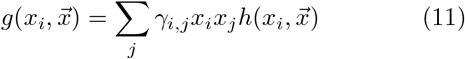
where *x_i_x_j_* is a mass action term *γ_i, j_* indicates the relative probability of interaction compared to mass action.Assuming the mass action hypothesis, the expected number of interactions is proportional to the product of the densities *x_i_* and *x_j_* of the pair of species. Other factors like the spatial component, consumer search efficiency,or handling time are aggregated in the preference matrix *γ* = [*γ_i, j_*]. This matrix measures pairwise interaction preferences: *γ_i, j_* >1 indicates the interaction is more likely to occur than expected, *γ_i, j_*<1 denotes a less favourable interaction and *γ_i, j_* = 1 is exactly the expectation based on mass action.

**Fig 5.**
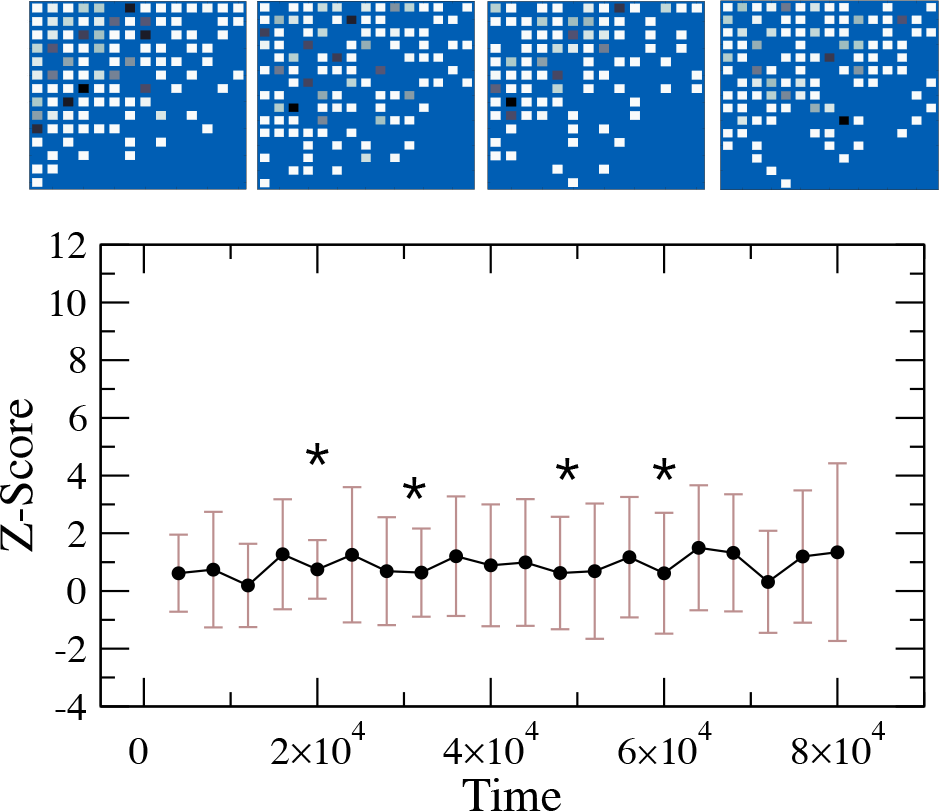
In the absence of space, the model does not tend to a pattern of significant nestedness (the average *Z* ≈ 0). The top row shows several snapshots of the host-phage interaction network taken at different evolutionary stages (whose location is given by stars in the curve). The binary structure of these matrices is nested but the quantitative preferences are found to be distributed in an anti-nested manner. The matrix at time *t* has been obtained by aggregating the interactions observed in the time period [*t* − Δ*t, t*] (here Δ*t* = 2000 time steps). In each matrix, darker colours represent higher preference of interactions. Tests for nestedness are based in the null model described in the main text.

When measuring nestedness in our system, we first adjust for the mass action effect (*x_i_x_j_*) to isolate interaction preferences. The incidence matrix *B* is related to the preference matrix by the following: *B_i, j_ = γ_i, j_x_i_x_j_* We compare the nestedness value in the preference matrix with an ensemble of random matrices having similar properties (Weitz et al. 2013; Beckett and Williams 2013). We use the null model proposed by (Staniczenko, Kopp, and Allesina 2013), which keeps the structural features of the network while swapping the order of weighted links (so-called ‘binary shuffle’). Specifically, the *Z*-score defines the statistical significance:

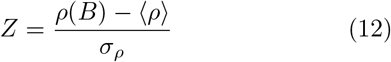
where 〈*ρ*〉 and *σ_ρ_* are the average value and the standard deviation of the network measure in a random ensemble, respectively. In this study, we will consider that host-phage interactions are significantly nested whenever the corresponding *Z* > 2 (*i.e., p* < 0:05 using the *Z*-test).

Figure 4 shows the statistical significance of nestedness in the *γ*-matrix tends to a high, well-defined value. In the absence of space, the same model does not tend to a pattern of significant nestedness (see Figure 5). Interestingly, the average connectance is also close to the reported value for the spatial simulation(*C* ≈ 0.45), and thus suggesting that nestedness is largely a consequence of spatial correlations associated to (transient) similarities between spatially close genomes.

## IV. Discussion

Host-virus ecological networks are characterised (among other things) by the presence of a nested organisation. Since nestedness has been proposed as a key attribute with a relevant role in community stability and diversity, to is specially important to understand its origins. Previous work using available host-phage networks has shown that nestedness appears to be a very common trait in most cases. What is less obvious is to determine the causal origins of this particular feature. A specially elegant work in this context is the study by (Beckett and Williams 2013) on the coevolutionary diversification of bacteria and phage using a lock-and-key model. This work aimed to account for both nestedness and modularity using a multi-strain chemostat system where the coevolving strains use a single resource. Genotypes are represented by means of a single scalar value, thus lacking our genotype space described by an explicit sequence hypercube. Importantly, the genetic matching is mediated by predefined functional correlations between “genotype distance” and key traits such as adsortion rates of phages on hosts.

In our analysis we have followed a rather different direction, by introducing a coevolution process where the specific choices of parameters (allowing populations to persist) is not relevant, genotypes are introduced in an explicit way and phenotypes are the same (as described by the kinetic parameters) for all genomes. In our study, the limited interactions among digital genomes associated to the presence of space play a key role in enhancing correlations and nestedness. We should expect that digital host genomes in a given neighborhood will also be relatively close among them through recent mutation events (in terms of Hamming distance) and exhibit closer ties with their parasites, which will also appear locally correlated. This necessarily helps enforcing the kind of correlations expected for nested graphs.

Despite its limitations, it is remarkable that such a simple set of assumptions recovers the nested organisation of these antagonistic systems. As it occurs with other relevant properties, spatial dynamics makes a difference when explicitly included in the description of ecosystem interactions. The loss of nestedness when global mixing is allowed clearly supports our conjecture. Future work should consider different extensions of our model, including spatial heterogeneity (which could lead to modularity) or theoretical developments that might help determine the validity and implications of our neutral approximation.

## Acknowledgments

The authors would like to thank the members of the Complex Systems Lab and our colleagues at the Santa Fe Institute for fruitful discussions. This work has been supported by the Botin Foundation by Banco Santander through its Santander Universities Global Division. This work was supported by the grants BFU2015-65037-P (S.F.E.) and FIS2016-77447-R (S.V.) from Spain Ministerio de Economía, Industria y Competitividad, AEI/MINEICO/FEDER and UE. The authors also thanks the Santa Fe Institute, where most of this work was done.

